# Cerebellar motor memory expression requires learned alterations to the activity of inhibitory molecular layer interneurons

**DOI:** 10.1101/2022.05.12.491667

**Authors:** Audrey Bonnan, Ke Zhang, Jason M. Christie

## Abstract

Procedural memories formed in the cerebellum in response to motor errors depend on changes to Purkinje cell (PC) spiking patterns that correct movement when the erroneous context is repeated. Because molecular layer interneurons (MLIs) inhibit PCs, learning-induced changes to MLI output may participate in reshaping PC spiking patterns. Yet, it remains unclear whether error-driven learning alters MLI activity and whether such changes are necessary for the memory engram. We addressed this knowledge gap by measuring and manipulating MLI activity in the flocculus of both sexes of mice before and after vestibulo-ocular reflex (VOR) adaptation. We found that MLIs are activated during vestibular stimuli and that their population response exhibits a phase shift after the instantiation of gain-increase VOR adaptation, a type of error-driven learning requiring climbing-fiber-mediated instructive signaling. Although acute optogenetic suppression of MLI activity did not affect baseline VOR performance, it negated the expression of gain-increase learning, demonstrating a specific causal role of MLI activity changes in motor memory expression. This effect was transitory; after a multi-day consolidation period, the expression of VOR gain-increase learning was no longer sensitive to MLI activity suppression. Together, our results indicate that error-driven alteration of MLI activity is necessary for labile, climbing-fiber-induced motor memory expression.

**Significance Statement:** In the cerebellum, motor learning induces an associative memory of the sensorimotor context of an erroneous movement that, when recalled, results in a new pattern of output that improves subsequent trials of performance. Our study shows that error-driven motor learning induces changes to the activity pattern of cerebellar molecular layer interneurons and that this new pattern of activity is required to express the corrective motor memory.

## Introduction

During procedural motor learning, the cerebellum stores memories of sensorimotor associations that, upon recall, modify movement (Bastian, 2006; Medina, 2011; Gao et al., 2012). It is well established that cerebellar Purkinje cells (PCs) adjust their simple spiking to include the development of phase shifts or well-timed pauses in firing coinciding with learned movements (Watanabe, 1984, 1985; Lisberger et al., 1994; Jirenhed et al., 2007; ten Brinke et al., 2015). Because PCs fire spontaneously at high rates *in vivo* (Thach, 1968) (~50–100 Hz), the loss or suppression of simple spiking releases premotor target neurons in the cerebellar nuclei from tonic inhibition, driving motor output (Heiney et al., 2014; Lee et al., 2015). Mossy fibers conveying sensory and motor information excite PCs via granule cell intermediaries, increasing PC simple spiking. Learning-induced synaptic plasticity triggered by the activity of climbing fibers weakens granule-cell-to-PC excitatory synaptic connections (Wang et al., 2014; Inoshita and Hirano, 2018). Such long-term depression (LTD) has emerged as an attractive cellular mechanism to explain acquired PC firing pauses in learned sensorimotor associations (Ito, 1982; Ito and Kano, 1982). However, transgenic mice deficient for cerebellar LTD continue to learn (Schonewille et al., 2011), indicating that additional or complementary mechanisms must also be necessary for the acquisition and expression of procedural motor memories (Boele et al., 2018).

In addition to excitation from granule cells, PCs also receive powerful inhibition from molecular layer interneurons (MLIs), which are similarly excited by granule cells. Thus, the net effect of sensorimotor stimulation on PC firing depends on the integration of both granule cell excitation and feedforward inhibition from MLIs (Mittmann et al., 2005; Mittmann and Hausser, 2007; Gaffield and Christie, 2017). Because MLI synapses are plastic (Kano et al., 1996; Jorntell and Ekerot, 2002, 2003; Rancillac and Crepel, 2004; Soler-Llavina and Sabatini, 2006; Kawaguchi and Hirano, 2007; Bender et al., 2009; Pugh and Jahr, 2011), error-driven reorganization of MLI output may contribute to learning-dependent changes to PC simple spiking patterns. In support of this idea, observations have shown cerebellar-dependent motor learning deficits in mice with PCs lacking GABA_A_ receptors (Wulff et al., 2009). Furthermore, associative learning alters MLI activity patterns (Ma et al., 2020). It remains unknown whether such changes in MLI activity are necessary for the expression of motor learning.

Here, we used a combination of *in vivo* functional recording and optogenetic activity perturbations to study the role of MLIs in adaptive oculomotor behavior. We discovered that MLI population dynamics exhibit reshaping during the acquisition of climbing-fiber-instructed learning and that, after this change, MLI activity becomes necessary for expression of the adapted behavioral response. Together, these findings contribute to a new understanding of local inhibitory circuits in cerebellar function: rather than simply acting as passive elements that determine PC excitability, MLIs undertake an active role in altering cerebellar output that corrects behavior, implicating their dynamics as a critical circuit feature underlying the motor memory engram.

## Materials and Methods

### Animals

All procedures were conducted at the Max Planck Florida Institute for Neuroscience on mature male and female mice (≥8 weeks for brain slice recording and ≥10 weeks for behavioral monitoring) using protocols approved by the Institutional Animal Care and Use Committee. Mice used in this study were heterozygous *kit::Cre* animals in which MLIs express Cre recombinase to permit their genetic targeting (Amat et al., 2017). The mice had *ad libitum* access to food and water and were normally held on a 12-hour light/dark cycle. However, where indicated, the mice were instead held in complete darkness for up to 5 days over the course of repeated training.

### Brain slice electrophysiology

For acute brain slice preparation, mice were anesthetized by intraperitoneal injection of ketamine/xylazine (100 and 10 mg/kg, respectively) and then transcardially perfused with cold saline (~4°C). After quickly removing the cerebellum by surgical dissection, parasagittal slices (200 μm) were sectioned from the vermis using a vibrating-blade microtome (VT1200S; Leica Biosystems) in an icy solution containing (in mM): 87 NaCl, 25 NaHO_3_, 2.5 KCl, 1.25 NaH_2_PO_4_, 7 MgCl_2_, 0.5 CaCl_2_, 10 glucose, and 75 sucrose. After sectioning, the slices were transferred to an incubation chamber containing a solution composed of (in mM): 128 NaCl, 26.2 NaHO_3_, 2.5 KCl, 1 NaH_2_PO_4_, 1.5 CaCl_2_, 1.5MgCl_2_ and 11 glucose. The slices were held in the incubation chamber for 40 min at 34°C and then at room temperature (23-25°C) thereafter until use. All solutions were oxygenated with carbogen gas (95% O_2_, 5% CO_2_) to equilibrium.

For experiments, the brain slices were placed in a submersion chamber under an upright microscope (BX51WI; Olympus) and continuously superfused with warmed bath solution (32-34°C). Recordings from MLIs were obtained in the cell-attached mode using gradient-contrast microscopy imaging for visualization. Recording pipettes (2-6 MΩ) were filled with a filtered solution containing (in mM): 119 NaCl, 2.5 KCl, 12 HEPES, 3 MgCl_2_, and 17 dextrose. Electrophysiological signals were measured using an amplifier and a digitizer (Multiclamp 700b and Digidata 1440A, respectively; Molecular Devices) controlled by commercial acquisition software (pClamp10; Molecular Devices). During electrophysiological recordings, light stimuli (λ565 nm) for optogenetics were delivered by wide-field epi-illumination from an unfiltered light-emitting diode (LED) (M565L3; Thorlabs). This light was launched through the back-port of the microscope and modulated (<1 kHz) with a current controller (LEDD1B; Thorlabs). All electrophysiological data were analyzed with Axograph X (Axograph).

### Surgical procedures

For viral injection surgeries, mice were anesthetized by continuous isoflurane gas (1–5%) and then placed on a stereotactic platform (Model 900, David Kopf Instruments) using ear bars. The anesthetic depth was determined by the absence of toe pinch responses. Continuous thermoregulation was provided by a heating plate with biofeedback to maintain physiological body temperature. A subcutaneous injection of lidocaine/bupivacaine was delivered to the scalp for local anesthesia, prior to a small incision being opened (< 2 mm) to allow for a craniotomy to be cut in the skull (< 0.5 µm in diameter). Through this opening, a glass micropipette containing AAV was advanced to the following coordinates (in mm from Bregma): X = ±2.35; Y = −5.65; Z = 3.2 - 3.4, and α = 10° for targeting the flocculus. For all injections, undiluted viral solution (titer ≥ 10^12^ vg/mL) was slowly infused into the target site (0.2–0.5 µL per injection). The micropipette was held in place (5–10 min) before withdrawal. Injected AAVs included AAV1.EF1α.DIO(*rev)*hChR2(H143R)-YFP (Penn Vector Core), AAV5.EF1α.DIO(*rev*)eNphr3.0-YFP (UNC Vector Core), and AAV1.CAG.DIO(*rev)*GCaMP6f (Penn Vector Core).

For the mice used in behavioral experiments, custom-made stainless steel headposts were installed during the surgery by removing a section of scalp on the center of the head and attaching the headpost to the exposed skull with dental cement (Metabond; Parkell). Optical fibers (200 µm, NA 0.22 with Ø 1.25 mm ferrules for optogenetics; 400 µm, NA 0.48 with Ø 1.25 mm ferrules for photometry; Thorlabs) were then implanted to target the flocculus (coordinates X = ± 3.35 mm; Y = 5.65 mm; α = −14°; *z* = 1.9 ± 0.1 mm). The optical fibers were secured in place using dental cement. In one mouse, a second small craniotomy was opened nearby to allow access for functional recording. This craniotomy was filled with silicon elastomer when not in use. All animals were allowed to recover after surgery under analgesia provided by injection of carprofen and buprenorphine SR-LAB (5 mg/kg and 0.35 mg/kg, respectively). After the onset of transgene expression (10–21 days), animals were used for behavioral monitoring or were sacrificed to harvest their brains for acute slice preparation.

### Behavioral monitoring, in vivo recording, and optogenetics

To measure VOR-evoked eye movements, mice were head-restrained on a rotating platform that was mounted on a rotary stage (T-RSW60C, Zaber Technology) that delivered horizontal sinusoidal vestibular stimuli (1 Hz). The left eye position was recorded using a machine-vision camera in response to these passive head rotations. We used commercial eye-tracking software (ETL-200; ISCAN) to record the pupil position (*P*) across time relative to the corneal reflection (*CR*) provided by an infrared LED mounted on the camera. Before each training session, we used a calibration procedure to estimate the radius of pupil rotation (*Rp*) based on the measured *P* relative to *CR* in response to a known camera rotation around the stage’s vertical axis (± 10°). A white-light-emitting LED was used to modulate pupil diameter by illuminating the eye at various light intensities. We then calculated the *Rp* for various pupil diameters according to: *Rp = Δ/sin (20°)*. This procedure allowed us to determine the angular eye position throughout an experiment using the formula: *Eye position (Ep) = arcsin [(P*_*1*_*-CR*_*1*_*)-(P*_*2*_*-CR*_*2*_*)/Rp]*. This value was then used to calculate the VOR gain as follows: *Gain = eye velocity/stage velocity*. This analysis was performed using custom-written MATLAB software (MathWorks).

Mice were trained for VOR learning by associating passive vestibular stimuli with moving visual stimuli (black and white bars) that were presented on two monitors placed in front of the animal. For gain-increase training, the vestibular stimulus was paired with a visual stimulus of the opposite direction (1.5x), whereas for gain-decrease training, the vestibular stimulus was paired with same-direction visual stimulation (0x). Sinusoidal stage movements (1 Hz; ± 5°; 20°/s peak velocity for gain-increase training or 1 Hz; ± 8.5°; 22°/s peak velocity for gain-decrease training) were controlled by custom-written software (LabVIEW, National Instruments), as were the visual stimuli, which moved relative to the rotation of the stage. Before training, mice were habituated to the behavioral set-up by being head-restrained for ~15 min on the platform for 1–2 days.

Baseline VOR performance was determined immediately before training in complete darkness. To limit darkness-induced pupil dilatation, pilocarpine (2% ophthalmic drops; Patterson Veterinary Supply) was applied (< 1 min) onto the eye after the calibration procedure. After training, the VOR was re-tested in darkness to measure learned changes in VOR performance. Measurements before and after training were obtained in the control condition as well as during optogenetic MLI activity suppression using 30 sec bouts of recording that were randomly interleaved. VOR gain changes were computed as the percentage difference relative to the control measurement obtained before training (ΔVOR). This normalization procedure facilitated comparison across sessions and between mice. The order of training sessions (gain-increase and gain-decrease) was randomized with at least two days between sessions. For multi-day gain-increase training, mice were removed from restraint after the end of each session and transferred to a light-free holding room for 24 h until the next training session; this was repeated over 5 consecutive days.

For optogenetic experiments during the VOR, MLI activity was suppressed using light pulses (λ589 nm; 1 pulse, 30 s) from a laser (CNI Optoelectronics Tech; MGL-F-589–200mW). For this, laser light was split into two lines with each line directed through independent acousto-optic modulators (AOMs) (AA Opto-Electronic; MTS110-A3-VIS) to control the laser power. From each AOM, laser light was launched into fiber ports (PAF-X-11-A; Thorlabs); patch cables then delivered light (8 mW out of patch cable) to the optical fiber implants targeting the flocculi of experimental animals. Eye displacement in response to the optogenetic activation or suppression of MLI activity was measured in head-restricted, quiescent mice using 240 ms pulses triggered every 10 s (λ473 nm or λ589 nm, respectively). The timing of eye movement onset was analyzed from ipsiversive stimulation recordings (1 pulse, 240 ms, 10 mW).

To record activity in an eNphr3.0-expressing mouse, a multi-electrode silicon probe (H6B; Cambridge NeuoTech) was slowly lowered into place in the flocculus through the exposed craniotomy. This was achieved using a manual manipulator targeting the same coordinates as the implanted optical fiber while the awake animal was held in head fixation. Three separate recording session were performed. Electrophysiological signals were acquired using an amplifier (RHD2132, Intan Technologies) readout on a controller interface and commercial software (Intan Technologies) at a sampling rate of 20 kHz. An implanted silver wire provided a ground signal. Laser light was delivered through the patch cable as a series of pulses (14 - 40 mW; 5 s; 0.1 Hz). The resulting data were automatically sorted using the Kilosort algorithm with manual curation performed using Phy2 software. Putative PC units were identified by their spiking properties and by the induced increase in firing when the flocculus was optogenetically disinhibited by MLI photo-suppression.

For photometry recordings, excitation light (λ470 nm; ~10–40 µW) from a fiber-coupled LED (M470F3; Thorlabs) was launched through a patch cable to the implanted optical fibers. Electrical signals from the custom-written control software triggered the current controllers (LEDD1B; Thorlabs) to toggle the LED on and off during the vestibular stimulus. The emitted fluorescence was collected through the same implanted optical fiber, passed back through the dichroic mirror mount, and detected using a femtowatt, visible wavelength photoreceiver (Model 2151; Newport) at a high sampling rate (2 kHz). Data points were averaged, producing an effective rate of activity measurement of 25 Hz. In interleaved trials, we used UV light (centered at λ405 nm; M405F1, Thorlabs) to excite GCaMP6f and record the isosbestic (calcium-insensitive) emission in response to vestibular stimulation. We used custom-written software to analyze these recordings (Python; Python Software Foundation). To quantify the timing and amplitude of the response, we fit the calcium activity waveform with a sinusoidal function and, from this fit, measured the phase of the peak activity relative to the stage position, as well as the peak-to-trough size of the evoked response (ΔF/F; with F defined as the baseline response in quiescence).

### Histology and fluorescence microscopy

For post-hoc examination of eNpHR3.0-YFP and GCaMP6f expression as well as confirmation of optical-fiber-placement, mice were deeply anesthetized by intraperitoneal injection of ketamine and xylazine (100 and 10 mg/kg, respectively) and then transcardially perfused with cold tris-buffered saline (TBS) followed by 4% paraformaldehyde (PFA) in TBS. After overnight post-fixation in PFA, the cerebellum was removed by dissection and cut into thin sections (60-80 µm) that were mounted onto glass slides for imaging on a confocal microscope (LSM 780; Zeiss) using the appropriate laser-excitation wavelengths and emission filter sets for each fluorophore.

### Statistical Analysis

All group values are presented as mean ± sem. The numbers of cells or animals used for each panel are indicated in the figures or the figure legends. Two-tailed paired t-test, repeated measures one-way analysis of variance (ANOVA) with Tukey’s multiple comparisons post-test, or repeated measures two-way ANOVA with Sidak’s multiple comparisons post-test were used to test for significant differences between two or more groups of data. For each data set, the statistical test used is indicated in the figure legends or in the text.

## Results

### Optogenetically induced MLI activity drives eye movements

The cerebellar flocculus is a site for learning-dependent oculomotor control, with PC activity regulating eye movement velocity and direction (Clopath et al., 2014; Payne et al., 2019). Therefore, to confirm that MLIs have the capacity to influence eye movements through their inhibitory effect on PCs, we applied an optogenetic approach to stimulate MLIs through implanted optical fibers targeting the flocculi of *kit::*Cre mice bilaterally injected with an adeno-associated virus (AAV) containing Cre-dependent ChR2 (*Figure 1A*). This targeting strategy has been previously shown to be effective in transducing MLIs with excitatory opsins (Amat et al., 2017). As assessed by videography of the left eye in head-restrained quiescent mice, photostimulating ChR2-expressing MLIs in either the contralateral or ipsilateral flocculus elicited eye movements whose trajectories included a combination of both horizontal and vertical components (*Figure 1B*).

**Figure 1.**
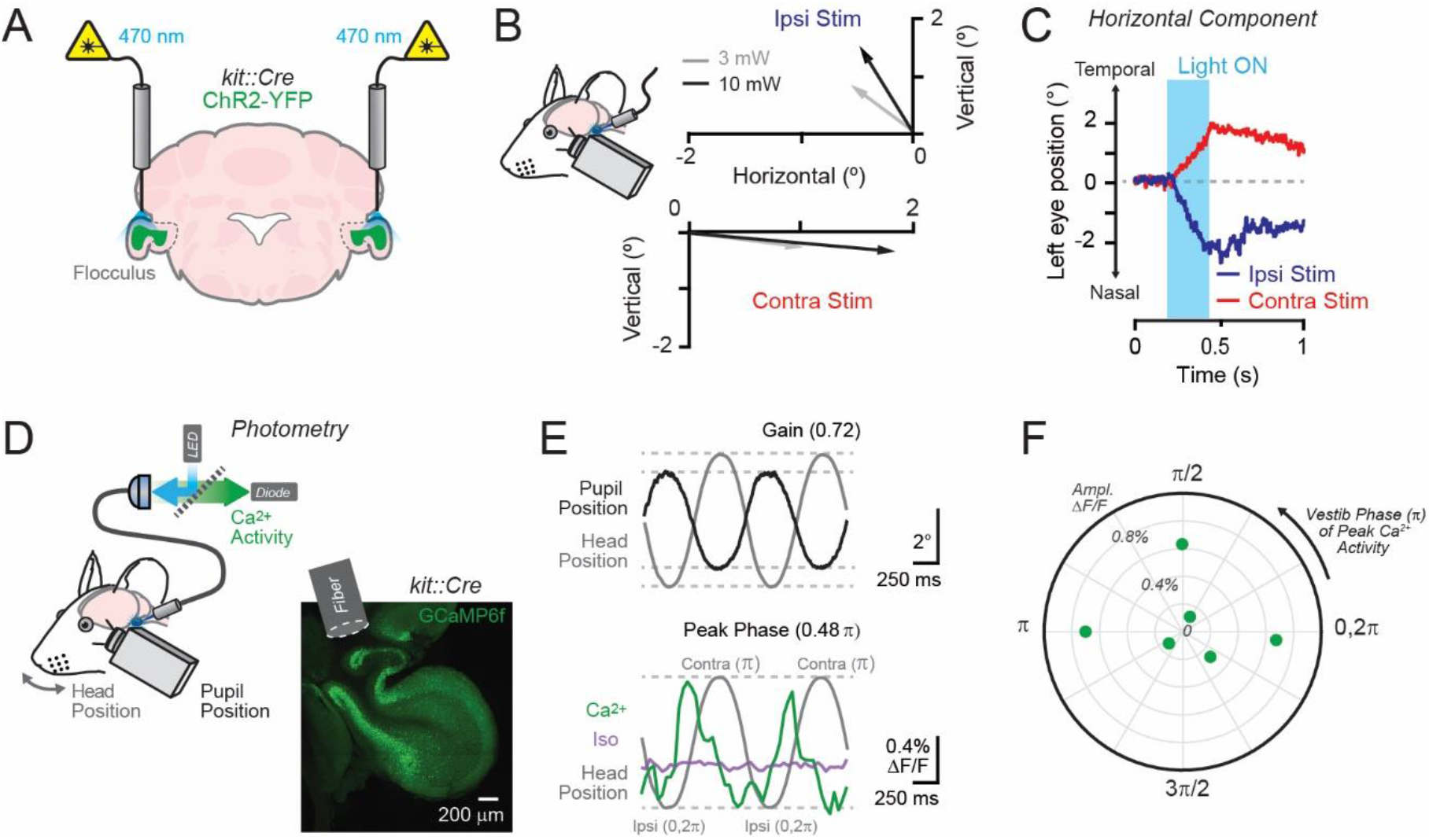
Optogenetic MLI activation evokes eye movement. **(A)** Mice expressing ChR2 in MLIs were bilaterally implanted with optical fibers targeting both flocculi. **(B)** Vector plot of the average direction and amplitude of left eye movements evoked by photostimulation of MLIs in the left (ipsiversive) or right (contraversive) flocculus at different laser powers (λ473 nm, 240 ms; n = 3 mice). **(C)** Decomposed horizontal eye movements from a quiescent mouse in response to unilateral floccular photostimulation of ChR2-expressing MLIs (10 mW). **(D)** For photometry, bulk GCaMP6f fluorescence was collected through an implanted optical fiber targeting the left flocculus as the VOR was passively elicited by sinusoidal vestibular stimulation (1 Hz) in darkness. GCaMP6f-expressing MLIs transduced using a Cre-dependent AAV are shown in the image with the approximate location of the optical fiber also indicated. **(E)** Top: trial-averaged VOR-evoked eye movements (black; head position in grey) in a GCaMP6f-expressing mouse. Bottom: calcium activity measurements from MLIs (green) with interleaved isosbestic measurements (purple) during the same recording. **(F)** The timing of peak calcium activity in the MLI population response relative to the phase of the vestibular stimulus for each mouse (each point represents an individual mouse; n = 6 mice total). The position along the radius corresponds to the peak amplitude of the calcium response.

Examining the isolated horizontal components, we observed that movement onset was well-timed to the light stimulus (response delay: 35.1 ± 3.7 ms; n = 3 mice). Movement velocity increased with the light intensity (3 mW: 7 ± 2.2°/s vs. 10 mW: 9.8 ± 2.1°/s, n = 3, p = 0.047, paired *t*-test; *Figure 1B*), indicating that motor kinematics were responsive to the level of MLI activation. The optogenetically evoked eye movements were directionally biased, dependent on the stimulus context. Activating MLIs in the contralateral flocculus moved the left eye away from the midline of the head. By contrast, stimulating MLIs in the ipsilateral flocculus moved the left eye toward the midline (*Figure 1B,C*). Thus, MLI activity can bidirectionally influence eye movements.

### MLIs are activated during the vestibulo-ocular reflex

Provided that optogenetically induced MLI activity is sufficient to drive eye movements, we sought to determine whether MLIs are intrinsically activated during the vestibulo-ocular reflex (VOR), a compensatory eye-movement-behavior generated contrary to head motion that helps maintain a stable gaze. To measure MLI population dynamics in the flocculus during the VOR, we employed fiber photometry. For this approach, we transduced MLIs in the left flocculus of *kit*::Cre mice with the calcium sensor GCaMP6f using a Cre-dependent AAV. We then recorded changes in bulk fluorescence using an implanted optical fiber while the VOR was elicited in darkness by passive sinusoidal head turns (*Figure 1D*).

The MLI population response modulated with the vestibular stimulus (*Figure 1E*), including activity peaks that exhibited a bias for specific phases of head motion. The phase in which population activity was the greatest varied depending on the individual animal. In some mice, MLI population activity peaked during the ipsiversive phase of vestibular motion (i.e., for MLIs in the left flocculus, activity peaked during leftward head turns) (*Figure 1F*, 4 out of 6 mice). In contrast, in other mice, MLI population activity peaked during the contraversive phase of vestibular motion (i.e., for the left flocculus, activity peaked during rightward head turns) (*Figure 1F*, 2 out of 6 mice). Because floccular granule cells spike in response to either ipsiversive or contraversive vestibular motion (Arenz et al., 2008), the phase-specific tuning that we observed in MLI population dynamics could reflect excitation driven by different granule-cell-input channels for each area sampled in our recordings. In summary, MLIs are engaged during VOR-evoked eye movements, with their population dynamics showing a continuum of phase-tuned responses throughout sinusoidal movements, similar to that observed for other oculomotor behaviors in mice (Badura et al., 2013).

### MLI activity is unnecessary for baseline VOR performance

To determine whether MLI activity contributes to shaping VOR-evoked eye movements, we used an optogenetic strategy to suppress their output. In our approach, we bilaterally injected Cre-dependent AAV containing the inhibitory opsin eNpHR3.0 (Gradinaru et al., 2010) into the flocculi of *kit::*Cre mice to transduce MLIs and implanted optical fibers targeting the infected regions (*Figure 2A*). We first confirmed that eNpHR3.0 activation suppressed MLI firing, even during prolonged periods of light exposure (30 s), using electrophysiological recordings from MLIs in acute cerebellar slices (*Figure 2B,C*). Extracellular electrophysiology measurements from putative PCs in an awake, eNpHR3.0-expressing mouse confirmed that MLI activity suppression disinhibited the flocculus when light pulses were delivered by an implanted optical fiber (*Figure 2D,E*).

**Figure 2.**
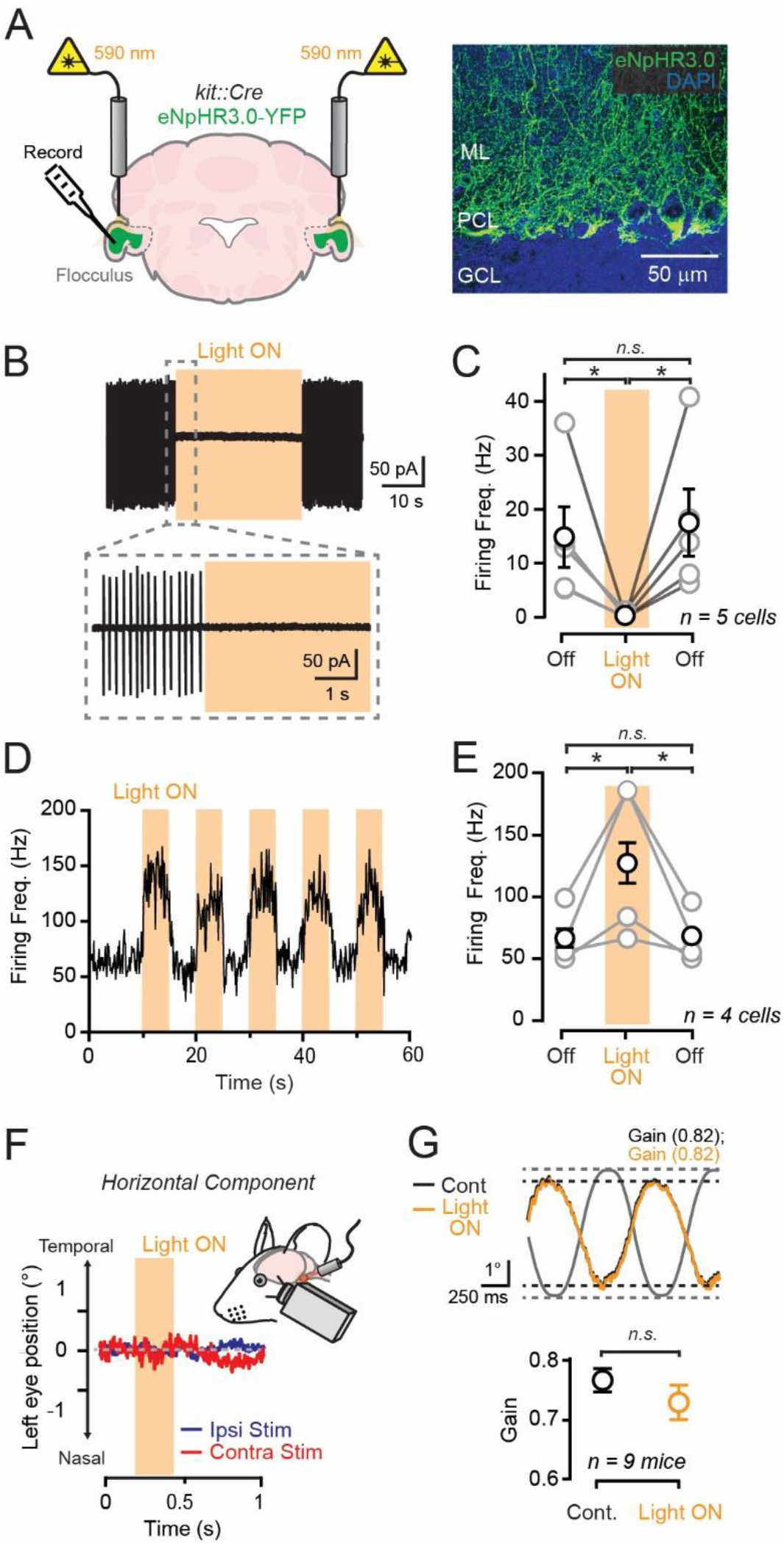
MLI activity suppression does not affect baseline VOR performance. **(A)** Light was delivered to eNpHR3.0-expressing MLIs using implanted optical fibers bilaterally targeting each flocculus. The image on the right shows YFP-tagged-eNpHR3.0 in MLIs from an example mouse. Molecular layer, ML; PC layer, PCL; granule cell layer, GCL. **(B)** Effect of eNpHR3.0 photoactivation (λ594 nm; 30 s, 3mW) on an MLI firing spontaneous spikes, measured in the cell-attached mode, in an acute cerebellar slice. **(C)** Summary plot showing the suppressive effect of eNpHR3.0 photoactivation on MLI firing (one-way ANOVA with Tukey’s multiple comparisons test; before vs. during, p = 0.0419, before vs. after, p = 0.8521, during vs. after, p = 0.0191). **(D)** The simple spiking of a putative floccular PC recorded in an awake eNpHR3.0-expressing mouse to optogenetic suppression of MLI activity (λ594 nm; 5 pulses at 0.1 Hz, 14 mW). **(E)** Summary plot showing the effect of MLI activity suppression on the spontaneous firing of putative PCs measured in three separate recording sessions (one-way ANOVA with Tukey’s multiple comparisons test; before vs. during, p = 0.0002, before vs. after, p = 0.9995, during vs. after, p = 0.0004). **(F)** Eye position measurements from a quiescent mouse as light pulses (λ590 nm, 240 ms, 8 mW) were delivered to the left (ipsiversive) or right (contraversive) flocculus to suppress MLI activity. **(G)** Top: average VOR-evoked eye movements in a mouse in the control condition (black) or during the bilateral suppression of MLI activity (orange). Light was delivered continuously to both flocculi for 30 s (λ590 nm; 8 mW) while the head was passively rotated in darkness using sinusoidal vestibular stimuli (1 Hz; head position in grey). Bottom: comparison of the VOR gain (the size of the evoked eye movement relative to the size of the vestibular stimulus) in the control condition and during optogenetic MLI activity suppression; not significant, s.; paired t-test, p = 0.067. All data are shown mean ± sem.

In head-fixed quiescent mice, brief (240 ms) optogenetic suppression of MLI activity in either the ipsiversive or contraversive flocculus failed to move the left eye (ipsiversive: Δ eye position during stimulus = 0.07 ± 0.08°, p = 0.46; contraversive: Δ eye position = 0.04 ± 0.02°, n = 3, p = 0.15, paired *t-*tests; *Figure 2F*). The absence of an evoked behavioral response suggests that spontaneous MLI activity is ineffective in regulating motor output outside of active behavior (Gaffield and Christie, 2017; Gaffield et al., 2018). We next suppressed MLI activity during the VOR by continuously delivering light to both flocculi for multiple trials of sinusoidal vestibular stimuli. However, this perturbation did not significantly affect the amplitude (gain) of the VOR (*Figure 2G*). This result indicates that although MLI activity is sufficient to impart eye movements and MLIs are activated during the VOR, the MLI output is dispensable for the VOR, at least in naïve animals with an already well-calibrated response.

### MLI activity is required for the expression of VOR learning

A poorly functioning VOR results in image instability during head motion. Therefore, the brain uses visual feedback to gauge how well vestibular stimuli are converted into eye movement coordinates. Retinal slip (visual motion) in the direction opposite to head motion indicates an under-performing VOR. These sensorimotor mismatches elicit climbing fiber activity in the flocculus that, through associative plasticity induction, changes the PC spiking pattern when the same vestibular context is repeated thereby increasing the VOR gain to restore distortion-free performance (Ito, 1982). We hypothesized that MLI activity may become necessary for VOR-evoked eye movements only after mice learn to re-calibrate their response to retinal slip errors. To test this idea, we trained mice with opposite-direction visual-vestibular-motion mismatches (1.5x; 60 min) that drove a learned increase in the VOR gain (ΔVOR 12.2 ± 2.0%, n = 9 mice, p = 0.0003, paired t-test). Bilateral optogenetic suppression of floccular MLI activity had a profound behavioral effect immediately after this learning. During the photo-suppression period, the adapted VOR response was fully negated, exhibiting a gain similar to that prior to learning (*Figure 3A,B*). The adapted response returned when MLI photo-suppression ceased. Light delivery to the flocculi of control *kit*::Cre mice injected with AAV containing Cre-dependent YFP did not influence the VOR gain, either before or after gain-increase learning (*Figure 3B*), confirming the specific effect of optogenetic MLI activity suppression on motor memory expression. Together, these results causally implicate the necessity of MLI activity in the expression of VOR gain-increase learning.

**Figure 3.**
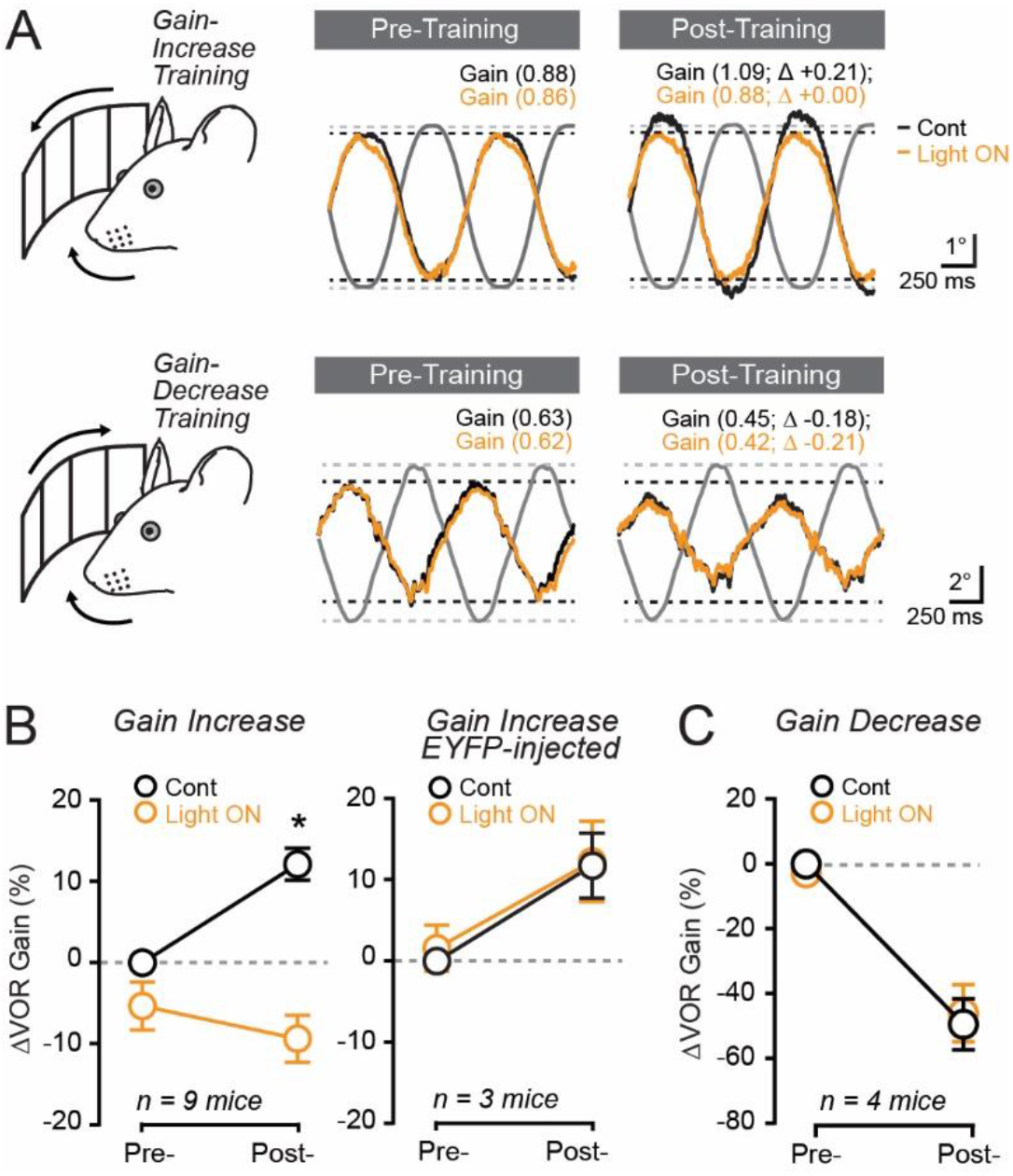
MLI activity suppression impairs the expression of gain-increase VOR learning. **(A)** Top: trial-averaged VOR-evoked eye movements in a mouse before (pre-) and after (post-) training with opposite-direction visual-vestibular motion mismatches (1.5x; 60 min). The responses were obtained in the control condition (black) and during optogenetic suppression of floccular MLI activity (orange). The black dotted lines indicate the amplitude of the baseline VOR in the control condition before training. Changes in gain (Δ) are a comparison to the baseline condition in the absence of optogenetic stimulation. Bottom: same mouse as above except for same-direction visual-vestibular motion mismatch training (0x; 60 min). **(B)** Left plot: summary showing the effect of MLI activity suppression on the expression of gain-increase learning (control vs. Light ON pre: p = 0.1315, control vs. Light ON post: p < 0.0001; two-way ANOVA with Sidak’s multiple comparisons). Right plot: the lack of effect of light delivery to mice expressing eYFP in MLIs before and after gain-increase training (control vs. Light ON pre: p = 0.71, control vs. Light ON post: p = 0.96; two-way ANOVA with Sidak’s multiple comparisons). **(C)** The effect of optogenetic MLI activity suppression on the expression of gain-decrease learning (control vs. Light ON pre: p = 0.90, control vs. Light ON post: p = 0.99; two-way ANOVA with Sidak’s multiple comparisons). Data are presented as mean ± sem.

Image instability can also elicit an adaptive decrease in the VOR gain when retinal slip occurs in the same direction as head motion. Because gain-decrease VOR learning is mechanistically distinct from gain-increase learning, and is instantiated independent of climbing-fiber-mediated signaling (Boyden et al., 2006; Kimpo et al., 2014; Bonnan et al., 2021), we tested whether MLI activity is also necessary for the expression of this form of cerebellar learning. In separate sessions, a cohort of the same mice were instead trained with same-direction visual-vestibular-motion mismatches (0x; 60 min) that resulted in a VOR gain decrease (52.7 ± 6.5% of baseline, n = 4 mice, p = 0.0039, paired t-test). Surprisingly, bilateral optogenetic suppression of floccular MLI activity had no effect on the gain of the adapted response (*Figure 3A,C*). We conclude that MLI activity is only involved in the expression of VOR learning induced by the activity of climbing fibers.

### MLI activity is reorganized during learning

The requirement for MLI activity in the expression of gain-increase learning is suggestive of a plasticity process that conditionally changes how MLIs respond during the VOR. Therefore, we used fiber photometry to measure the dynamics of MLI population activity before and after training mice with opposite-direction visual-vestibular-motion mismatches (1.5x; 60 min). With the acquisition of gain-increase learning, the behavior-evoked MLI activity pattern exhibited a profound reorganization. For each mouse, the peak population response shifted from one preferred phase of vestibular motion to another (*Figure 4A,B*). Generally, MLI populations that displayed an initial bias for ipsiversive vestibular motion became responsive to contraversive motion whereas MLI populations initially responsive to contraversive vestibular motion became responsive to ipsiversive motion (phase difference post-training: 0.74 ± 0.08 π rad; n = 5 mice; p = 0.0008, paired *t*-test; *Figure 4A,B*). We observed an exception in one mouse in which the initial peak in activity was not lost; however, a second peak developed timed to the opposite phase of vestibular motion. Across the mice, the calcium response amplitude did not change after learning (peak ΔF/F: 0.38 ± 0.12% and 0.47 ± 0.17% pre-training and post-training, respectively, p = 0.63, paired *t*-test), suggesting that the overall level of their population activity remained the same. Together, these results indicate that the response patterns of floccular MLIs are not fixed during the VOR; rather, their activity is subject to phase reversals during gain-increase learning.

**Figure 4.**
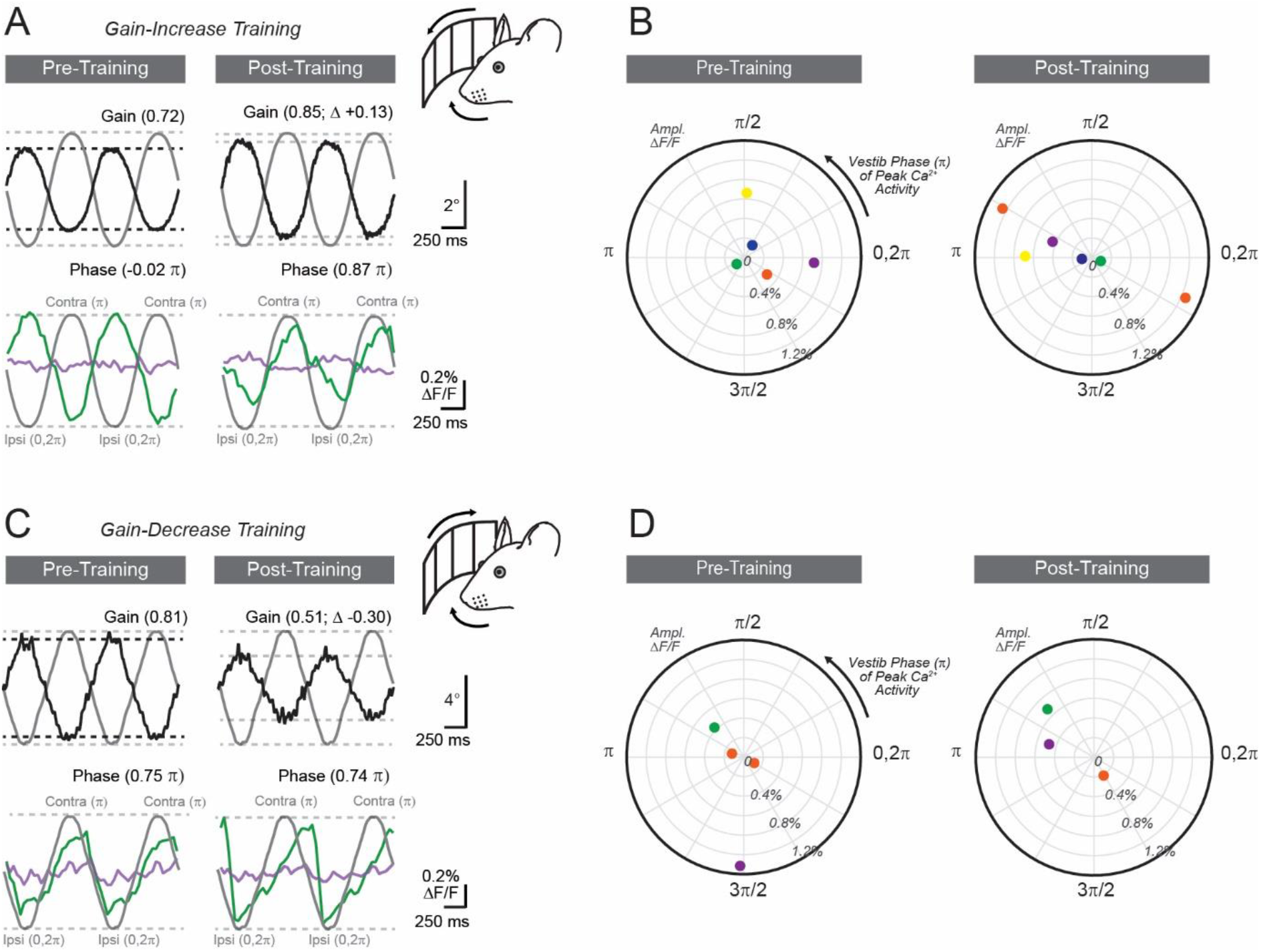
MLI population activity is restructured in response to gain-increase VOR learning. **(A)** VOR-evoked eye movements and the simultaneously acquired population activity of MLIs, measured in the left flocculus of a GCaMP6f-expressing mouse using fiber-photometry, in trials before and after gain-increase learning. **(B)** Summary plots showing the timing of peaks in the MLI population response, relative to the phase of the vestibular stimulus, in mice before (left) and after (right) gain-increase learning. Individual mice, represented as single points, are color coded. Note, one mouse (red) developed a second peak in calcium activity after learning. **(C**,**D)** Same as panels A and B but for gain-decrease VOR learning. Color-coding corresponds to the same mice in panel B.

In separate sessions, we performed MLI activity measurements in some of the same mice as they were trained with same-direction visual-vestibular-motion mismatches (0x; 60 min). Despite the acquisition of gain-decrease learning, the preferred phase of the MLI population response (phase difference post-training: −0.25 ± 0.17 π rad, n = 3 mice, p = 0.27, paired *t*-test; *Figure 4C,D*) and amplitude (peak ΔF/F: 0.91 ± 0.64% and 0.46 ± 0.13% pre-training and post-training, respectively, p = 0.54, paired *t*-test) were relatively stable. Therefore, the requirement for MLI activity in motor memory expression developed in accord with learned changes to the structure of their population response. This result suggests that the dependence of motor memory expression on MLI activity likely results from an alteration of MLI population dynamics. We conclude that phase shifts in MLI activity during VOR gain-increase learning are a correlate of climbing-fiber-induced motor memory expression.

### MLI-mediated inhibition is transiently required for motor memory expression

Prior research has led to the hypothesis that labile motor memories initially triggered in the cerebellar cortex are slowly transferred to downstream premotor locations for lasting storage (Miles and Lisberger, 1981; Lisberger, 2021). Based on this model of cerebellar function, we reasoned that the expression of gain-increase learning should only transiently depend on neural activity in the flocculus, up until the point of long-term consolidation (Kassardjian et al., 2005; Anzai et al., 2010). To examine the temporal requirement of MLI activity for the expression of climbing-fiber-mediated oculomotor learning, we trained mice over five consecutive daily sessions with opposite-direction visual-vestibular motion mismatches (1.5x; 60 min/day) and assessed the effect of optogenetic MLI activity suppression on VOR performance before and after each training session (*Figure 5A*). Mice were housed in darkness between each session to prevent reversal learning and to allow for memory consolidation.

**Figure 5.**
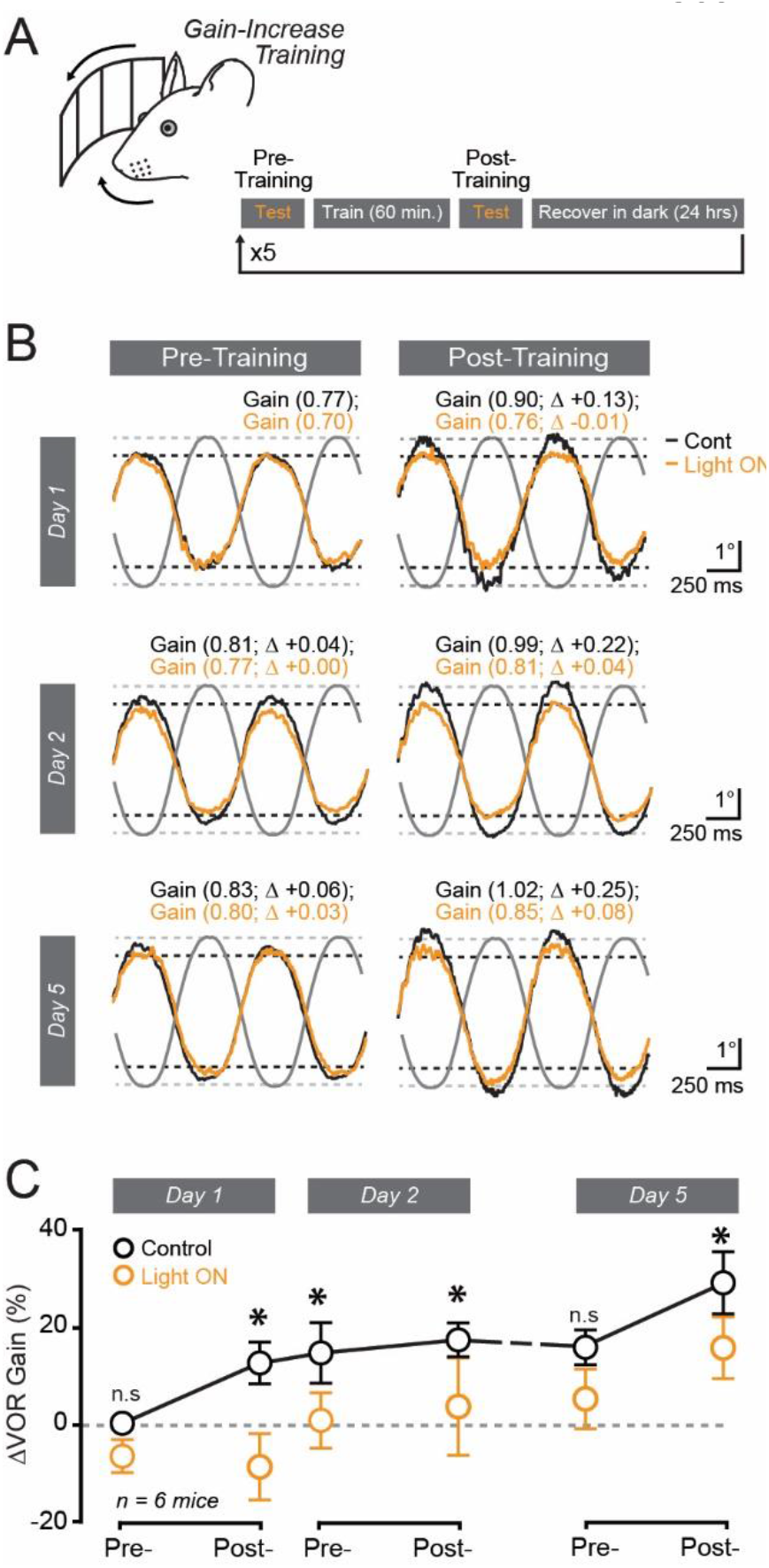
Transient requirement for MLI activity in the expression of gain-increase VOR learning. **(A)** Mice received five consecutive days of opposite-direction visual-vestibular-motion mismatch training. At the start and end of each session, the VOR was measured in the control condition and during optogenetic MLI activity suppression. Mice were held in a light-free environment between sessions. **(B)** Trial-averaged VOR eye movements from the same mouse measured on different training days. Responses obtained in the control condition (black) and during MLI activity suppression (orange) are superimposed. Black dotted line indicates the amplitude of the baseline VOR response measured on the first day of training. Changes in gain (Δ) are a comparison to the initial baseline condition in the absence of optogenetic stimulation. **(C)** Plot showing the effect of MLI activity suppression on the VOR gain for the progression of multi-day gain-increase learning across mice. Statistics for control vs. Light ON: Day1 Pre and Post, p = 0.9993 and p <0.0001, respectively; Day2 Pre and Post: p = 0.0073 and p = 0.0212, respectively; Day5 Pre and Post, p = 0.8411 and p = 0.0470, respectively; two-way ANOVA with Sidak’s post comparisons. Data are presented as mean ± sem.

In the control condition, the gain-increase learning acquired during the first training session was apparent at the start of the second session, indicating that the mice had retained a memory of the previous day’s learning (ΔVOR: 12.6 ± 4.4% and 14.8 ± 6.3%, respectively, n = 6 mice, p = 0.797, paired t-test; *Figure 5B,C*). Unlike the first session, optogenetic MLI activity suppression had a dramatic effect on the VOR at the start of the second session, reducing the gain to the baseline level measured the day before (*Figure 5B,C*). Thus, expression of the remanent motor memory still required MLI activity. The optogenetic perturbation had a similar negating effect on motor memory expression at the end of the second session, suggesting that further training did not obviate the role of MLI activity in the adapted response (*Figure 5B,C*).

The mice continued to exhibit an adapted VOR gain at the start of the fifth session, relative to the baseline response prior to training (ΔVOR: 15.9 ± 3.7%, n = 6 mice, p = 0.002, paired t-test; *Figure 5B,C*), highlighting the long-term retention of the previously acquired learning. Interestingly, MLI activity suppression at the start of this last session failed to affect the VOR gain. Thus, after several days of training, lasting motor memory expression no longer required MLI activity, a result supporting a consolidation process outside of the cerebellar cortex. During this fifth session, opposite-direction visual-vestibular-motion mismatch training further increased the VOR gain (ΔVOR: 13.5 ± 3.8% over the course of the session, n = 6 mice, p = 0.016, paired t-test), suggesting that the new learning compounded on the previously consolidated memory. Notably, optogenetic MLI activity suppression fully negated the expression of this new learning as, during light delivery, the VOR gain matched the level at the start of the session (*Figure 5B,C*). We conclude that this newly formed climbing-fiber-mediated motor memory requires MLI activity.

## Discussion

We demonstrate a causative role for MLIs in corrective motor memory expression that depends on error-driven restructuring of behavior-evoked MLI population dynamics. By delineating the importance of local inhibitory interneurons in motor memory engrams, our results contribute new insight into the neural-circuit mechanisms underlying procedural cerebellar learning.

### Phasic modulation of MLI population activity during the VOR

Our functional recordings indicate that floccular MLIs are engaged during the VOR and structure their population dynamics to modulate during different phases of vestibular motion. Across measurements from different mice, MLI activity encompassed the entirety of reflexive bidirectional eye movements evoked by sinusoidal vestibular stimuli. Thus, MLIs are activated across multiple types of motor behaviors (Badura et al., 2013; Jelitai et al., 2016; Astorga et al., 2017; Chen et al., 2017; Gaffield and Christie, 2017). We suspect that the spectrum of phase-tuned MLI population responses that we observed among mice is attributable to random sampling that captured activity from different MLI ensembles, perhaps driven by granule cell inputs that preferentially fire to either ipsiversive or contraversive head turns (Arenz et al., 2008). Such a compartmental organization could support functional modules within the flocculus that may be important for controlling distinct agonist and antagonist muscle groups of the eyes (Van der Steen et al., 1994; Voogd and Wylie, 2004). Our use of photometry did not provide the spatial resolution necessary to determine how individual cells in the flocculus modulate their activity during the VOR, including both increases and decreases in output that compose the population response, precluding a more precise determination of MLI coding features.

As in a prior study (Heiney et al., 2014), we found that suppressing spontaneous PC simple spiking through optogenetically induced MLI stimulation elicits motor action. The resulting eye movement velocity was responsive to the MLI activity level, and the movement direction depended on whether MLIs in either the ipsiversive or contraversive flocculus were activated. Floccular PCs activation also elicits eye movements, but in a direction opposite to that evoked by MLI stimulation (Nguyen-Vu et al., 2013; Voges et al., 2017; Bonnan et al., 2021). Thus, the positive and negative modulation of PC firing, driven by granule cell excitation or MLI inhibition, respectively, can produce a contrasting range of movements with different kinematic and directional features. Despite their engagement during the behavior, we found that MLIs do not contribute to shaping the gain of VOR eye movements in naïve mice, a result consistent with prior observations showing that acutely blocking floccular activity does not affect the performance of an already well-calibrated response (Nagao and Kitazawa, 2003; Kassardjian et al., 2005). Together, these results indicate that although the activity of the vestibulo-cerebellum tracks the VOR performance, including the participation of MLIs, it does not always exert online control of the behavior.

### MLI activity is necessary for motor memory expression

The vestibular nuclei, the final node of the brainstem loop controlling the VOR, integrate direct excitation from mossy fibers as well as inhibition from PCs. Because the relative timing of these two inputs determines the output of the vestibular nuclei during oculomotor behavior, learning-induced phase shifts in PC simple spike activity (Watanabe, 1984, 1985; Lisberger et al., 1994) enables the cerebellum to conditionally influence eye movement velocity (Ito, 1982). Plastic re-weighting of granule cell-PC synapses has long been viewed as the cellular substrate for producing changes in PC firing during memory recall, with LTD and long-term potentiation (LTP) being putative mechanisms for learned increases and decreases in VOR gain, respectively (Titley and Hansel, 2016). We discovered that MLI activity is also necessary for the expression of gain-increase VOR learning, indicating the participation of MLIs in sculpting PC spiking dynamics underlying motor memory engrams. Prior studies using pharmacology or constitutive knockout mice to disinhibit the cerebellum have also suggested a role for molecular layer inhibition in motor learning (Wulff et al., 2009; Tanaka et al., 2013). Unlike the use of optogenetics in our study, these approaches lack the specificity and reversibility to disambiguate the role of MLIs in regulating the induction of VOR learning (Rowan et al., 2018) from the role of MLIs in motor memory expression.

Because MLIs modulate PC simple spiking activity through their inhibitory activity (Brown et al., 2019), changes in inhibition could contribute to learned patterns of PC output. In this way, pauses in PC firing that develop with learning might be best explained by an acquired increase in inhibitory output by MLIs timed to the adapted response (ten Brinke et al., 2015). Our results do not preclude the possibility that learning-induced MLI plasticity occurs in combination with parallel fiber-PC LTD, or other sites of plasticity in the cerebellar cortex, to synergistically manifest learned behavior during memory recall (Gao et al., 2012; Boele et al., 2018). Further work will be necessary to determine whether loss of PC simple spiking during learning is sensitive to MLI activity suppression or if it is attributable to a process independent of MLI-mediated inhibition (Johansson et al., 2014). The different requirements for MLI activity in the expression of gain-increase versus gain-decrease VOR learning further emphasizes that these different learning types rely on distinguishable plasticity mechanisms (Boyden et al., 2004). In summary, multiple memory pathways are available in the cerebellum to compose engrams that support disparate learned behaviors that may include either the strengthening or weakening of motor output, the former requiring MLI activity.

### Learning induces restructuring of MLI population activity

We found that MLI population dynamics undergo a profound reorganization during gain-increase VOR learning, fully reversing the preferred phase of their activation in response to sinusoidal vestibular motion. Learning-induced alterations of MLI activity have also been observed in mice performing a reward-driven odor discrimination task (Ma et al., 2020). Therefore, experience-dependent alteration of MLI activity occurs across a range of learned behaviors. That said, MLI population activity did not change over the course of gain-decrease VOR learning. Thus, the dependence of motor memory expression on MLI activity develops in accord with the induced alterations to their population dynamics. Because gain-increase but not gain-decrease VOR learning results from climbing-fiber-mediated signaling (Boyden et al., 2006; Nguyen-Vu et al., 2013; Kimpo et al., 2014; Rowan et al., 2018; Bonnan et al., 2021), this result suggests a role for climbing fibers in instructing the restructuring of MLI population activity.

Learning-induced changes in MLI population activity could result from plasticity at their granule cell inputs (Jorntell and Ekerot, 2003; Rancillac and Crepel, 2004; Soler-Llavina and Sabatini, 2006; Bender et al., 2009). For example, granule-cell-mediated excitation of MLIs potentiates when stimulated in conjunction with the activity of climbing fibers (Jorntell and Ekerot, 2003). This LTP would increase the level of feedforward MLI-mediated inhibition of PCs transforming the PC firing pattern to granule cell input, which could ultimately alter behavior. We posit that learning-induced phase shifts in the MLI population dynamics modify the timing of PC-mediated inhibition of the vestibular nuclei, relative to mossy-fiber-mediated excitation, such that the same vestibular signal results in a larger eye movement after learning. Based on our observation of directionally biased movement during unilateral floccular stimulation, the addition of contraversive MLI activity would drive responses away from the midline, contributing to larger amplitude eye movements during the VOR increasing the response gain. In our activity measurements, however, learning-induced changes included both gains and losses of contraversive- and ipsiversive-motion-preferring MLI population responses. It is possible that phase-tuned MLI ensembles have different roles in neural circuit function. Thus, the symmetrical loss and gain of both contraversive- and ipsiversive-motion-preferring MLI population activity could have different behavioral effects.

### Transient requirement for MLI activity in motor memory expression

Our findings show that MLIs are only transiently involved in motor memory engrams because, during the recall of lasting memories of previously acquired gain-increase learning, optogenetic MLI activity suppression had no effect on the adapted VOR response. These results are congruent with previous investigations (Kassardjian et al., 2005; Anzai et al., 2010) showing that acute pharmacological block of floccular activity prevents the expression of VOR gain-increase adaptation immediately after the training procedure but not after a multi-day consolidation period (i.e., > 72 hours). Taken together, our results support the conclusion that motor memories requiring MLI activity first form in the cerebellar cortex. However, as these memories are transferred over time to the cerebellar nuclei for long-term retention (Mauk, 1997; Lisberger, 2021), the role of MLIs in the engram is obviated. We did not determine if and/or how MLI activity may be reshaped during the consolidation process, but MLI activity dynamics are most likely conducive for further restructuring because new gain-increase learning requiring MLI activity can be acutely acquired after the consolidation process.

Our study emphasizes the importance of MLI activity restructuring in the expression of labile climbing-fiber-induced motor engrams, thus adding an extra dimension to the role of MLIs in behaviorally relevant computations (Jorntell et al., 2010; Kim and Augustine, 2021). Beyond the cerebellum, our results also contribute to the general understanding that local inhibitory circuits play a critical role in organizing neural activity underlying memory engrams throughout the brain (Li et al., 2013; Courtin et al., 2014; Cummings and Clem, 2020).

## Acknowledgements

We thank Samantha Amat for laboratory assistance and the GENIE program (Janelia Research Campus, including Drs. Jayaraman, Kerr, Kim, Looger, and Svoboda) for freely providing GCaMP6f to the neuroscience community. This work was supported by National Institutes of Health Grants NS112289 and NS105958 (J.M.C.), as well as the Max Planck Florida Institute for Neuroscience.

## References

Amat SB, Rowan MJM, Gaffield MA, Bonnan A, Kikuchi C, Taniguchi H, Christie JM (2017) Using c-kit to genetically target cerebellar molecular layer interneurons in adult mice. PloS one 12:e0179347.

Anzai M, Kitazawa H, Nagao S (2010) Effects of reversible pharmacological shutdown of cerebellar flocculus on the memory of long-term horizontal vestibulo-ocular reflex adaptation in monkeys. Neuroscience research 68:191–198.

Arenz A, Silver RA, Schaefer AT, Margrie TW (2008) The contribution of single synapses to sensory representation in vivo. Science (New York, NY) 321:977–980.

Astorga G, Li D, Therreau L, Kassa M, Marty A, Llano I (2017) Concerted Interneuron Activity in the Cerebellar Molecular Layer During Rhythmic Oromotor Behaviors. The Journal of neuroscience : the official journal of the Society for Neuroscience 37:11455–11468.

Badura A, Schonewille M, Voges K, Galliano E, Renier N, Gao Z, Witter L, Hoebeek FE, Chedotal A, De Zeeuw CI (2013) Climbing fiber input shapes reciprocity of Purkinje cell firing. Neuron 78:700–713.

Bastian AJ (2006) Learning to predict the future: the cerebellum adapts feedforward movement control. Current opinion in neurobiology 16:645–649.

Bender VA, Pugh JR, Jahr CE (2009) Presynaptically expressed long-term potentiation increases multivesicular release at parallel fiber synapses. The Journal of neuroscience : the official journal of the Society for Neuroscience 29:10974–10978.

Boele HJ, Peter S, Ten Brinke MM, Verdonschot L, ACH IJ, Rizopoulos D, Gao Z, Koekkoek SKE, De Zeeuw CI (2018) Impact of parallel fiber to Purkinje cell long-term depression is unmasked in absence of inhibitory input. Science advances 4:eaas9426.

Bonnan A, Rowan MMJ, Baker CA, Bolton MM, Christie JM (2021) Autonomous Purkinje cell activation instructs bidirectional motor learning through evoked dendritic calcium signaling. Nature communications 12:2153.

Boyden ES, Katoh A, Raymond JL (2004) Cerebellum-dependent learning: the role of multiple plasticity mechanisms. Annual review of neuroscience 27:581–609.

Boyden ES, Katoh A, Pyle JL, Chatila TA, Tsien RW, Raymond JL (2006) Selective engagement of plasticity mechanisms for motor memory storage. Neuron 51:823–834.

Brown AM, Arancillo M, Lin T, Catt DR, Zhou J, Lackey EP, Stay TL, Zuo Z, White JJ, Sillitoe RV (2019) Molecular layer interneurons shape the spike activity of cerebellar Purkinje cells. Scientific reports 9:1742.

Chen S, Augustine GJ, Chadderton P (2017) Serial processing of kinematic signals by cerebellar circuitry during voluntary whisking. Nature communications 8:232.

Clopath C, Badura A, De Zeeuw CI, Brunel N (2014) A cerebellar learning model of vestibulo-ocular reflex adaptation in wild-type and mutant mice. The Journal of neuroscience : the official journal of the Society for Neuroscience 34:7203–7215.

Courtin J, Chaudun F, Rozeske RR, Karalis N, Gonzalez-Campo C, Wurtz H, Abdi A, Baufreton J, Bienvenu TC, Herry C (2014) Prefrontal parvalbumin interneurons shape neuronal activity to drive fear expression. Nature 505:92–96.

Cummings KA, Clem RL (2020) Prefrontal somatostatin interneurons encode fear memory. Nature neuroscience 23:61–74.

Gaffield MA, Christie JM (2017) Movement Rate Is Encoded and Influenced by Widespread, Coherent Activity of Cerebellar Molecular Layer Interneurons. The Journal of neuroscience : the official journal of the Society for Neuroscience 37:4751–4765.

Gaffield MA, Rowan MJM, Amat SB, Hirai H, Christie JM (2018) Inhibition gates supralinear Ca(2+) signaling in Purkinje cell dendrites during practiced movements. eLife 7.

Gao Z, van Beugen BJ, De Zeeuw CI (2012) Distributed synergistic plasticity and cerebellar learning. Nature reviews Neuroscience 13:619–635.

Gradinaru V, Zhang F, Ramakrishnan C, Mattis J, Prakash R, Diester I, Goshen I, Thompson KR, Deisseroth K (2010) Molecular and cellular approaches for diversifying and extending optogenetics. Cell 141:154–165.

Heiney SA, Kim J, Augustine GJ, Medina JF (2014) Precise control of movement kinematics by optogenetic inhibition of Purkinje cell activity. The Journal of neuroscience : the official journal of the Society for Neuroscience 34:2321–2330.

Inoshita T, Hirano T (2018) Occurrence of long-term depression in the cerebellar flocculus during adaptation of optokinetic response. eLife 7.

Ito M (1982) Cerebellar control of the vestibulo-ocular reflex--around the flocculus hypothesis. Annual review of neuroscience 5:275–296.

Ito M, Kano M (1982) Long-lasting depression of parallel fiber-Purkinje cell transmission induced by conjunctive stimulation of parallel fibers and climbing fibers in the cerebellar cortex. Neuroscience letters 33:253–258.

Jelitai M, Puggioni P, Ishikawa T, Rinaldi A, Duguid I (2016) Dendritic excitation-inhibition balance shapes cerebellar output during motor behaviour. Nature communications 7:13722.

Jirenhed DA, Bengtsson F, Hesslow G (2007) Acquisition, extinction, and reacquisition of a cerebellar cortical memory trace. The Journal of neuroscience : the official journal of the Society for Neuroscience 27:2493–2502.

Johansson F, Jirenhed DA, Rasmussen A, Zucca R, Hesslow G (2014) Memory trace and timing mechanism localized to cerebellar Purkinje cells. Proceedings of the National Academy of Sciences of the United States of America 111:14930–14934.

Jorntell H, Ekerot CF (2002) Reciprocal bidirectional plasticity of parallel fiber receptive fields in cerebellar Purkinje cells and their afferent interneurons. Neuron 34:797–806.

Jorntell H, Ekerot CF (2003) Receptive field plasticity profoundly alters the cutaneous parallel fiber synaptic input to cerebellar interneurons in vivo. The Journal of neuroscience : the official journal of the Society for Neuroscience 23:9620–9631.

Jorntell H, Bengtsson F, Schonewille M, De Zeeuw CI (2010) Cerebellar molecular layer interneurons-computational properties and roles in learning. Trends in neurosciences 33:524–532.

Kano M, Kano M, Fukunaga K, Konnerth A (1996) Ca(2+)-induced rebound potentiation of gamma-aminobutyric acid-mediated currents requires activation of Ca2+/calmodulin-dependent kinase II. Proceedings of the National Academy of Sciences of the United States of America 93:13351–13356.

Kassardjian CD, Tan YF, Chung JY, Heskin R, Peterson MJ, Broussard DM (2005) The site of a motor memory shifts with consolidation. The Journal of neuroscience : the official journal of the Society for Neuroscience 25:7979–7985.

Kawaguchi SY, Hirano T (2007) Sustained structural change of GABA(A) receptor-associated protein underlies long-term potentiation at inhibitory synapses on a cerebellar Purkinje neuron. The Journal of neuroscience : the official journal of the Society for Neuroscience 27:6788–6799.

Kim J, Augustine GJ (2021) Molecular Layer Interneurons: Key Elements of Cerebellar Network Computation and Behavior. Neuroscience 462:22–35.

Kimpo RR, Rinaldi JM, Kim CK, Payne HL, Raymond JL (2014) Gating of neural error signals during motor learning. eLife 3:e02076.

Lee KH, Mathews PJ, Reeves AM, Choe KY, Jami SA, Serrano RE, Otis TS (2015) Circuit mechanisms underlying motor memory formation in the cerebellum. Neuron 86:529–540.

Li H, Penzo MA, Taniguchi H, Kopec CD, Huang ZJ, Li B (2013) Experience-dependent modification of a central amygdala fear circuit. Nature neuroscience 16:332–339.

Lisberger SG (2021) The Rules of Cerebellar Learning: Around the Ito Hypothesis. Neuroscience 462:175–190.

Lisberger SG, Pavelko TA, Bronte-Stewart HM, Stone LS (1994) Neural basis for motor learning in the vestibuloocular reflex of primates. II. Changes in the responses of horizontal gaze velocity Purkinje cells in the cerebellar flocculus and ventral paraflocculus. Journal of neurophysiology 72:954–973.

Ma M, Futia GL, de Souza FMS, Ozbay BN, Llano I, Gibson EA, Restrepo D (2020) Molecular layer interneurons in the cerebellum encode for valence in associative learning. Nature communications 11:4217.

Mauk MD (1997) Roles of cerebellar cortex and nuclei in motor learning: contradictions or clues? Neuron 18:343–346.

Medina JF (2011) The multiple roles of Purkinje cells in sensori-motor calibration: to predict, teach and command. Current opinion in neurobiology 21:616–622.

Miles FA, Lisberger SG (1981) Plasticity in the vestibulo-ocular reflex: a new hypothesis. Annual review of neuroscience 4:273–299.

Mittmann W, Hausser M (2007) Linking synaptic plasticity and spike output at excitatory and inhibitory synapses onto cerebellar Purkinje cells. The Journal of neuroscience : the official journal of the Society for Neuroscience 27:5559–5570.

Mittmann W, Koch U, Hausser M (2005) Feed-forward inhibition shapes the spike output of cerebellar Purkinje cells. The Journal of physiology 563:369–378.

Nagao S, Kitazawa H (2003) Effects of reversible shutdown of the monkey flocculus on the retention of adaptation of the horizontal vestibulo-ocular reflex. Neuroscience 118:563–570.

Nguyen-Vu TD, Kimpo RR, Rinaldi JM, Kohli A, Zeng H, Deisseroth K, Raymond JL (2013) Cerebellar Purkinje cell activity drives motor learning. Nature neuroscience 16:1734–1736.

Payne HL, French RL, Guo CC, Nguyen-Vu TB, Manninen T, Raymond JL (2019) Cerebellar Purkinje cells control eye movements with a rapid rate code that is invariant to spike irregularity. eLife 8.

Pugh JR, Jahr CE (2011) NMDA receptor agonists fail to alter release from cerebellar basket cells. The Journal of neuroscience : the official journal of the Society for Neuroscience 31:16550–16555.

Rancillac A, Crepel F (2004) Synapses between parallel fibres and stellate cells express long-term changes in synaptic efficacy in rat cerebellum. The Journal of physiology 554:707–720.

Rowan MJM, Bonnan A, Zhang K, Amat SB, Kikuchi C, Taniguchi H, Augustine GJ, Christie JM (2018) Graded Control of Climbing-Fiber-Mediated Plasticity and Learning by Inhibition in the Cerebellum. Neuron 99:999–1015.e1016.

Schonewille M, Gao Z, Boele HJ, Veloz MF, Amerika WE, Simek AA, De Jeu MT, Steinberg JP, Takamiya K, Hoebeek FE, Linden DJ, Huganir RL, De Zeeuw CI (2011) Reevaluating the role of LTD in cerebellar motor learning. Neuron 70:43–50.

Soler-Llavina GJ, Sabatini BL (2006) Synapse-specific plasticity and compartmentalized signaling in cerebellar stellate cells. Nature neuroscience 9:798–806.

Tanaka S, Kawaguchi SY, Shioi G, Hirano T (2013) Long-term potentiation of inhibitory synaptic transmission onto cerebellar Purkinje neurons contributes to adaptation of vestibulo-ocular reflex. The Journal of neuroscience : the official journal of the Society for Neuroscience 33:17209–17220.

ten Brinke MM, Boele HJ, Spanke JK, Potters JW, Kornysheva K, Wulff P, Ac IJ, Koekkoek SK, De Zeeuw CI (2015) Evolving Models of Pavlovian Conditioning: Cerebellar Cortical Dynamics in Awake Behaving Mice. Cell reports 13:1977–1988.

Thach WT (1968) Discharge of Purkinje and cerebellar nuclear neurons during rapidly alternating arm movements in the monkey. Journal of neurophysiology 31:785–797.

Titley HK, Hansel C (2016) Asymmetries in Cerebellar Plasticity and Motor Learning. Cerebellum (London, England) 15:87–92.

Van der Steen J, Simpson JI, Tan J (1994) Functional and anatomic organization of three-dimensional eye movements in rabbit cerebellar flocculus. Journal of neurophysiology 72:31–46.

Voges K, Wu B, Post L, Schonewille M, De Zeeuw CI (2017) Mechanisms underlying vestibulo-cerebellar motor learning in mice depend on movement direction. The Journal of physiology 595:5301–5326.

Voogd J, Wylie DR (2004) Functional and anatomical organization of floccular zones: a preserved feature in vertebrates. The Journal of comparative neurology 470:107–112.

Wang W, Nakadate K, Masugi-Tokita M, Shutoh F, Aziz W, Tarusawa E, Lorincz A, Molnár E, Kesaf S, Li YQ, Fukazawa Y, Nagao S, Shigemoto R (2014) Distinct cerebellar engrams in short-term and long-term motor learning. Proceedings of the National Academy of Sciences of the United States of America 111:E188–193.

Watanabe E (1984) Neuronal events correlated with long-term adaptation of the horizontal vestibulo-ocular reflex in the primate flocculus. Brain research 297:169–174.

Watanabe E (1985) Role of the primate flocculus in adaptation of the vestibulo-ocular reflex. Neuroscience research 3:20–38.

Wulff P, Schonewille M, Renzi M, Viltono L, Sassoe-Pognetto M, Badura A, Gao Z, Hoebeek FE, van Dorp S, Wisden W, Farrant M, De Zeeuw CI (2009) Synaptic inhibition of Purkinje cells mediates consolidation of vestibulo-cerebellar motor learning. Nature neuroscience 12:1042–1049.

